# Utah Array Characterization and Histological Analysis of a Multi-Year Implant in Non-Human Primate Motor and Sensory Cortices

**DOI:** 10.1101/2022.08.27.505114

**Authors:** Paras R. Patel, Elissa J. Welle, Joseph G. Letner, Hao Shen, Autumn J. Bullard, Ciara M. Caldwell, Alexis Vega-Medina, Julianna M. Richie, Hope E. Thayer, Parag G. Patil, Dawen Cai, Cynthia A. Chestek

## Abstract

The Utah array is widely used in both clinical studies and neuroscience. It has a strong track record of safety. However, it is also known that implanted electrodes promote the formation of scar tissue in the immediate vicinity of the electrodes, which negatively impacts the ability to record neural waveforms. This scarring response has been primarily studied in rats and mice, which may have a very different response than primate brain. Here, we present a rare nonhuman primate histological dataset (n=1 rhesus macaque) obtained 848 and 590 days after implantation in two brain hemispheres. For 2 of 4 arrays that remained within the cortex, NeuN was used to stain for neuron somata at 3 different electrode depths. Images were filtered and denoised, with neurons then counted in the vicinity of the arrays as well as a nearby section of control tissue. Additionally, 3 of 4 arrays were imaged with a scanning electrode microscope (SEM) to evaluate any materials damage that might be present. Overall, we found a 63% percent reduction in the number of neurons surrounding the electrode compared to control areas. In terms of materials, the arrays remained largely intact with metal and Parylene C present, though tip breakage and cracks were observed on many electrodes. Overall, these results suggest that the tissue response in the nonhuman primate brain shows similar neuron loss to previous studies using rodents. Electrode improvements, for example using smaller or softer probes, may therefore substantially increase the neuronal recording yield in primate cortex.

## 1. Introduction

Brain-machine interfaces (BMIs) offer patients living with motor or sensory impairments – often resulting from injuries to the spinal cord, nerves or muscles – the chance to regain movement or restore sensation [1]–[3]. State-of-the-art BMIs require invasive technology that rests on direct electrical connection between healthy neural tissue and an external computer. The first clinical BMI trial began in 2004 and showed spinal cord injury patient Matthew Nagle able to operate a computer cursor and open and close a robotic hand [4]. Increasingly advanced medical feats occurred in the intervening years: patients have independently fed themselves a drink [5], felt the touch of a loved one [6], [7], controlled the movement of their own physically reanimated arm [8], regained touch-pressure sensation [9] and fist-bumped a former U.S. president [10]. The expanding list of BMI capabilities are underscored by engineering breakthroughs, such as sophisticated decoding algorithms that extract useful information from noisy data or low-power amplifiers that analyze brain signals with a fraction of the computational bandwidth previously needed [11], [12].

Yet, one aspect of BMIs has remained constant: the neural device responsible for interfacing directly with the brain’s cortex called the Utah Electrode Array (UEA), first published in the late 1990s [13]. The UEA features a 4 mm x 4 mm silicon body composed of 100 silicon shanks, each 1.5 mm long, arranged in a 10 × 10 grid. Electrodes are located at the tip of each shank and the grid spacing ensures each electrode is spaced 400 µm apart [14]. The array of 100 individual electrodes is implanted into the motor or sensory cortices of the brain. There, each electrode records electrical signals sent between neurons and sends the information to an external computer for real time data processing [15]. The UEA is the only intracortical device approved for clinical BMI use [16], a status that cemented its architecture for over three decades.

Despite the longest running clinical BMI study lasting approximately 5.4 years, the majority of BMIs studies report a worsening of two key measurements within months to one year after UEA implant: a decrease in recorded amplitude of electrical signals from individual or small groups of neurons, and a decrease in the total number of working electrodes on the UEA [17] [18]–[20]. Poor quality neural signals and low numbers of working electrodes limit the progress of BMIs towards the goal of replicating natural, high-precision movements and sensory inputs for BMI users [21].

There may be several reasons for the limited quality and quantity of neural signals recorded on UEAs. Studies of long-term UEA implants in brain tissue report fewer neurons and more tissue scarring and inflammation near electrodes [22]–[28]. In 2005, Biran, Martin, and Tresco published observations of the density of neurons around silicon shank electrodes, which are similar to Utah array shanks in size and material, and determined a decrease in neural density extending roughly 200 µm from the surface of a silicon shank [29]. In UEAs, this distance would affect the entire recording region between two adjacent electrodes that sit 400 µm apart [30]. In addition to possible decreases in neuronal density, the UEA electrodes appear to degrade under the constant exposure to the warm, watery, and high-salinity environment in the brain [31]. Reactive oxides – which are linked to the degradation of electrical devices [32] – were found in elevated numbers inside the scar region that was shown to form around UEA electrodes [33]. Extensive scar tissue and resulting encapsulation of UEAs in peripheral nerves were found to lead to Parylene C delamination, cracking, and thinning, as well as cracking of the conductive electrode coating [31], [34].

Much of the data on UEA degradation and the foreign body response was gathered in feline or rodent models [23], [28], [35]–[37], leaving the obvious question of what occurs during human clinical trials. A recent study on explanted UEAs after 0.5 and 2.7 years in the cortex of two BMI patients looked at the signal quality and material degradation, and found greater tissue encapsulation and worsening electrode coating degradation exhibited on longer implants [38]. Similarly, histopathology of tissue surrounding two UEAs in one patient confirmed similar scarring and widespread necrosis seen previously in animal models that correlated with signal degradation [39]. However, clinical trials are rare as is the histological analysis of human brain tissue. Non-human primates (NHPs) are often used to test advanced BMIs as a substitute to human patients.

One group led by John Donoghue examined changes in both electrode material and neural tissue histology after long-term UEA implantation in the cortex of over two dozen NHPs [18]. In this study, approximately 80% of UEAs failed completely while implanted, the majority of which failed within the first year of implant [18]. Explanted UEAs showed overall signs of material fatigue and degradation, while histological analysis of the neural tissue showed substantial presence of inflammatory markers and decrease in neuron density near the electrode holes [40]. These extensive studies laid the groundwork for understanding why and how UEAs fail in long-term brain implantation. Yet, a study quantifying changes in neuron density near electrode holes coupled with a visual analysis of the electrodes’ material degradation is still needed.

Here, we add to the collective knowledge of changes in neuronal density and UEA integrity after multi-year implantation in a rare dataset from the cortex of one NHP. We analyze the neuron density surrounding UEAs explant sites in the motor and sensory cortices in both the left and right hemispheres after 1.6 and 2.3 years, respectively. The neuron density around electrode holes is compared to nearby non-implanted tissue to quantify the change induced by UEA presence. We expand upon previous preliminary analysis of neuron density counted manually by introducing a semi-automated counting methodology applied to all electrode holes [41]. We also analyze images of the UEAs using scanning electron microscopy (SEM) to determine changes in the electrode surface and the spatial arrangement of degradation within the array. This study furthers our understanding of long-term UEA implantation in the cortex of an NHP, moving the field one step closer to understanding the implication of clinical BMI use.

## 2. Methods

### 2.1 Experimental set-up

This study examines the histological tissue response and material changes of four UEAs (Blackrock Microsystems, Salt Lake City, UT) implanted in the sensory and motor cortices of a single rhesus macaque NHP (Figure 1). The NHP was involved in a BMI study and trained for brain control tasks of individual finger movements [42], [43]. The NHP was also involved in corticocortical processing experiments using ketamine [44]. The NHP was euthanized at the termination of these experimental timelines due to successful completion of experimental objectives coupled with deteriorating health. The UEAs were extracted, cleaned, imaged, and analyzed for material changes. The brain tissue under the UEAs was sliced, stained, imaged, and analyzed for changes in neuron density.

**Figure 1.**
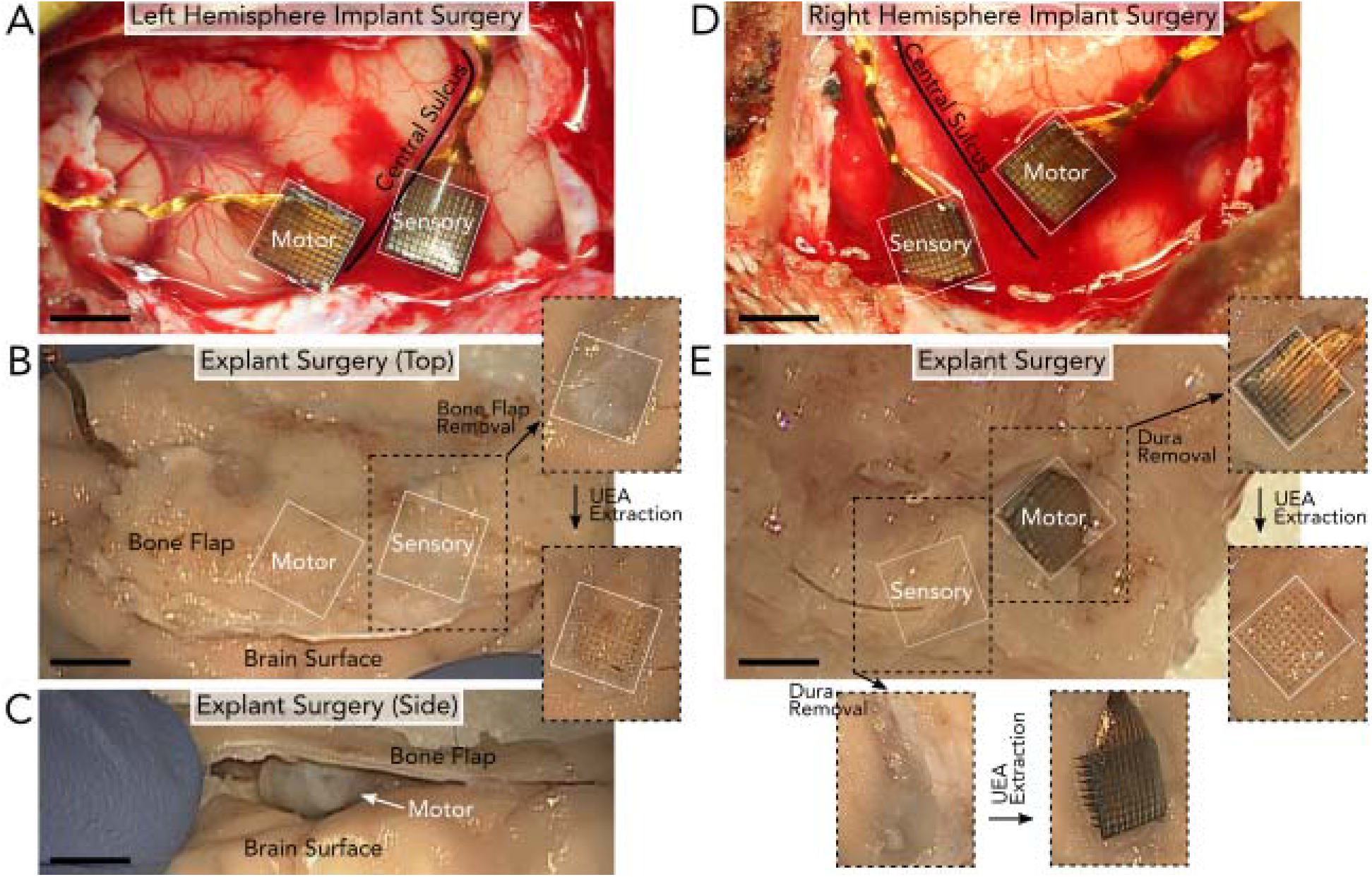
Surgical implantation and explantation of UEAs after 848 days in the left hemisphere (A-C) and 590 days in the right hemisphere (D-E) of the NHP. A) Left hemisphere implantation of two UEAs in the motor and sensory cortices on either side of the central sulcus. B) Explantation of the UEAs in (A) involved removing the section of bone (bone flap) above the arrays. After the bone flap and dura were removed, the UEAs could be extracted. Removing the left sensory UEA revealed clear holes in the tissue. C) However, the UEA in the left motor cortex was fully encapsulated by tissue and no longer implanted in the brain surface, as seen from the image taken of the side of the tissue. D) Right hemisphere implantation of two UEAs in the motor and sensory cortices on either side of the central sulcus. E) Explantation of the UEAs in (D) involved removing the bone flap above the arrays to reveal the two UEAs. After the bone flap was removed, the UEAs could be extracted. Both UEAs were partially or fully implanted in the tissue at the time of explant. All scale bars are 4mm.

### 2.2 UEA implantation

All animal procedures were approved by the University of Michigan Institutional Animal Care & Use Committee. Two UEAs were implanted in the primary motor (M1) and sensory (S1) cortex of each hemisphere. The left hemisphere was implanted on August 20, 2015 (Figure 1A-C) and the right hemisphere on May 4, 2016 (Figure 1D, E). Each UEA was of standard architecture: 100 electrodes at the tips of 1.5 mm-long shanks and a 6 cm length wire bundle. The electrode tips of three UEAs were coated with iridium oxide and implanted in the left motor (LM), left sensory (LS), and right sensory (RS) cortices. The right motor cortex (RM) was implanted with a UEA with platinum-coated electrode tips. The UEAs in left and right sensory cortices were fabricated with an experimental aluminum oxide coating prior to Parylene C insulation [45].

The NHP was placed in a stereotaxic frame after induction with general anesthesia during each implantation surgery. The location of the craniotomy over the central sulcus was estimated using the stereotaxic setup and a craniotomy and durotomy were performed over the region of implant. The UEAs were manually positioned and then impacted into the cortical tissue using a pneumatic inserter (Blackrock Microsystems, Salt Lake City, UT), seen in Figure 1A and 1D. The dura was closed over the UEAs and sealed with PRECLUDE Pericardial Membrane (Gore, Flagstaff, AZ) and DuraGen (Integra LifeSciences, Princeton, NJ). The bone flap was replaced and fastened with titanium bone screws (DePuy Synthes, Paoli, PA). Silicone elastomer (Kwik-Cast, World Precision Instruments, Sarasota, FL) and dental acrylic (A-M Systems, Sequim, WA) were applied to secure the wire bundles to the skull.

### 2.3 UEA and brain tissue extraction

All four UEAs were extracted on December 15, 2017, after 848 days of implantation in the left hemisphere (Figure 1B, C) and 590 days of implantation in the right hemisphere (Figure 1E). The terminal surgical extraction protocol required that perfusion not be performed while the NHP was under anesthesia. Therefore, post-mortem perfusion began approximately 4 minutes after death, as confirmed by veterinary staff. The NHP was anesthetized with ketamine and then sacrificed with euthanasia solution (VetOne, Boise, ID). The NHP was then transcardially perfused with heparinized (10 U/mL) 1X phosphate buffered saline (PBS) solution (BP3994, Fisher, Waltham, MA) until the exudate was clear, followed by approximately 1 L of 4% (w/v) paraformaldehyde (PFA) fixative (19208, Electron Microscopy Sciences, Hatfield, PA) in 1X PBS.

After perfusion, the dental acrylic and overlying bone flap were removed with a handheld drill. Dural growth on top of the UEAs was removed. The brain sections containing the UEAs were excised and placed in 4% PFA for 72 hours at 4 °C (Figure 1B, E). At this point the UEAs were removed with fine forceps and immediately placed in a chemical disinfectant (Benz-All, Xttrium Laboratories, Inc., Mount Prospect, IL) overnight. UEAs were switched to 1X PBS after approximately 24 hours to be preserved for future analysis. After UEA extraction, the brain sections were returned to 4% PFA for an additional 48 hours at 4 °C and then stored in 1X PBS at 4 °C.

### 2.4 Tissue slicing

Brain sections were trimmed of excess tissue and the implant portions were separated from each other. The implant portions were placed in 4% PFA for 5 days followed by 8 days in 1X PBS at 4 °C. Implant portions were cryoprotected in 30% sucrose (S0389, Sigma Aldrich, St. Louis, MO) in 1X PBS at 4 °C for 26 days and then frozen at -80 °C in optimal cutting temperature compound (Tissue-Tek, Sakura Finetek USA, Inc., Torrance, CA). The tissue was sliced perpendicular to the implantation sites in 100 µm thick sections at -16 °C on a cryostat. Tissue slices were stored in 0.02% sodium azide (DSS24080, Dot Scientific Inc., Burton, MI) in 1X PBS at 4 °C until immunohistochemical labeling. Throughout this study slices are referred to by their final depth in hundreds of microns from the surface of the brain. For example, slice 13 contains tissue 1200-1300 µm from the top of the implant portion.

### 2.5 Tissue staining

Slices at varying depths down the UEA shank were selected for tissue staining. Tissue slices were blocked and permeabilized with a mixture of StartingBlock PBS Blocking Buffer (37538, Thermo Scientific, Waltham, MA) and 1% Triton X-100 (9002-93-1, Sigma Aldrich, St. Louis, MO) overnight at 4 °C followed by three 30-minute washes in 1X PBS containing 0.5% Triton X-100, referred to as 0.5% PBST, at room temperature. The tissue was incubated with primary antibody at a 1:250 dilution in 0.5% PBST with 0.02% sodium azide for 48 hours at 4 °C. The following primary antibody was used to stain for neurons, mouse anti-neuronal nuclei (NeuN, MAB377, MilliporeSigma, Burlington, MA). Primary antibody incubation was followed by three 30-minute washes in 0.5% PBST at room temperature. The tissue was incubated in secondary antibody at a 1:250 dilution in 0.5% PBST with 0.02% sodium azide for 24 hours at 4 °C. The following secondary antibody was used, anti-mouse Alexa Fluor 647 (715-605-150, Jackson ImmunoResearch Laboratories, Inc., Carlsbad, CA). Finally, the tissue slices were washed in room temperature 0.5% PBST two times at two-hour intervals and kept in 1X PBS overnight. All slices were stored at 4 °C in 1X PBS with 0.02% sodium azide until imaged.

### 2.6 Tissue imaging

Tissue slices were imaged on a Zeiss LSM 780 Confocal Microscope (Carl Zeiss AG, Oberkochen, Germany) with a 20X objective. Images were collected with an approximately 0.4-0.6 µm X and Y pixel size and 2 µm z-step for the total 100 µm depth of the slice. The NeuN stain was imaged at a wavelength of 633 nm (NeuN). Laser intensity was adjusted manually to prevent pixel saturation, corrected in the Z direction to also prevent saturation, and ranged from 1.2-80% laser power. The gain and contrast were altered during image processing in ImageJ.

### 2.7 Tissue analysis

Neuron density around electrode sites was calculated from the NeuN tissue images and compared to the neuron density in non-implanted tissue. Viable non-implant tissue sites were selected in areas outside regions of visible damage. Electrode sites and non-implant tissue regions were cropped to 400 µm by 400 µm sections in MATLAB (Mathworks, Natick, MA) centering on the electrode hole. The image depth was cropped to the center 40 or 70 µm of the 100 µm-thick slice. The total 3D volume was 6.4×10^6^ µm^3^ or 11.2×10^6^ µm^3^.

Each cropped image was pre-processed in ImageJ (U. S. National Institutes of Health, Bethesda, Maryland). The pixel intensity was normalized across z-stacks to the highest signal-to-noise ratio z-stack using the histogram matching feature [46]. The image was filtered with a mean 50-pixel filter and the background was subtracted to remove pixelated noise. The image was then denoised with a 3D Gaussian 2-pixel radius filter to remove individual pixels with abnormally high intensity.

Then, the pre-processed image was read into MATLAB for 3D visualization using the Volume Viewer application. A 3D view of the neurons was generated using the isosurface feature with a unique isosurface value for each slice, as determined by a trained operator to accurately match the original image. A 2D image of the slice was imported back into ImageJ for cell counting. The image was smoothed and converted to 8-bit grayscale. The range of particle sizes used to identify neurons was determined by a trained operator measuring the smallest and largest neurons. The Analyze Particles program was run to locate neurons and a trained operator reviewed the resulting identifications for misidentified or unidentified neurons. The neuron density was calculated by dividing the total neuron count by the total volume.

### 2.8 UEA scanning electron microscopy imaging

To determine if there was any electrode degradation, UEAs were cleaned and imaged. First, UEAs were removed from 1X PBS and soaked in deionized water for 1 hour to detach any remaining brain tissue. UEAs were air dried for 1 hour prior to affixation to SEM stubs (16111, Ted Pella, Redding, CA) with carbon tape (16073, Ted Pella, Redding, CA). UEAs were imaged in a TESCAN Rise SEM (Tescan Orsay Holding, Brno–Kohoutovice, Czech Republic) at 20 kV using the low vacuum secondary detector. UEAs were tilted to approximately 20 degrees for maximum visibility of electrode tips. Images were collected of the whole array and of each quadrant of 5×5 electrode shanks. Backscatter mode images were also collected to detect cracks in the Parylene C insulation.

### 2.9 Electrode analysis

Images of each UEA quadrant of 5×5 electrode shanks were analyzed for six categories of degradation: electrode tip breakage (TB), cracks in metal electrode coating (CC), below-electrode tip shank fracture (SF), unidentified or abnormal debris (AB), Parylene C cracks (PC) and Parylene C peeling or delamination (PD). Examples of each category are shown in Figure 3B as identified on LS (Figure 3A), except for Parylene C delamination which was found on RM. Three trained operators scored each shank as exhibiting (1) or not exhibiting (0) the degradation category. Scores were averaged across operators and then rounded to 0 or 1. The outer three rows of electrode shanks were statistically compared to the inner 4×4 electrode shanks in an ANOVA test (alpha < 0.05). Analysis was conducted using MATLAB.

## 3. Results

### 3.1 Analysis of neuron density

We analyzed the tissue slices found under the LS array at three depths along the length of electrode shanks at 800-900 µm, 1000-1100 µm, and 1200-1300 µm and under the RS array at 1600-1700 µm, 1700-1800 µm, and 2000-2100 µm. Figure 2 depicts the electrode (Figure 2B) and non-implanted tissue (Figure 2C) regions of interest analyzed in slice 11 for the LS array (Figure 2A). Representative images of the three main stages of the analysis are shown in Figure 2B and 2C: the original image (top), the image after filtering and processing (middle), and the image analyzed with Analyze Particles in ImageJ (bottom).

**Figure 2.**
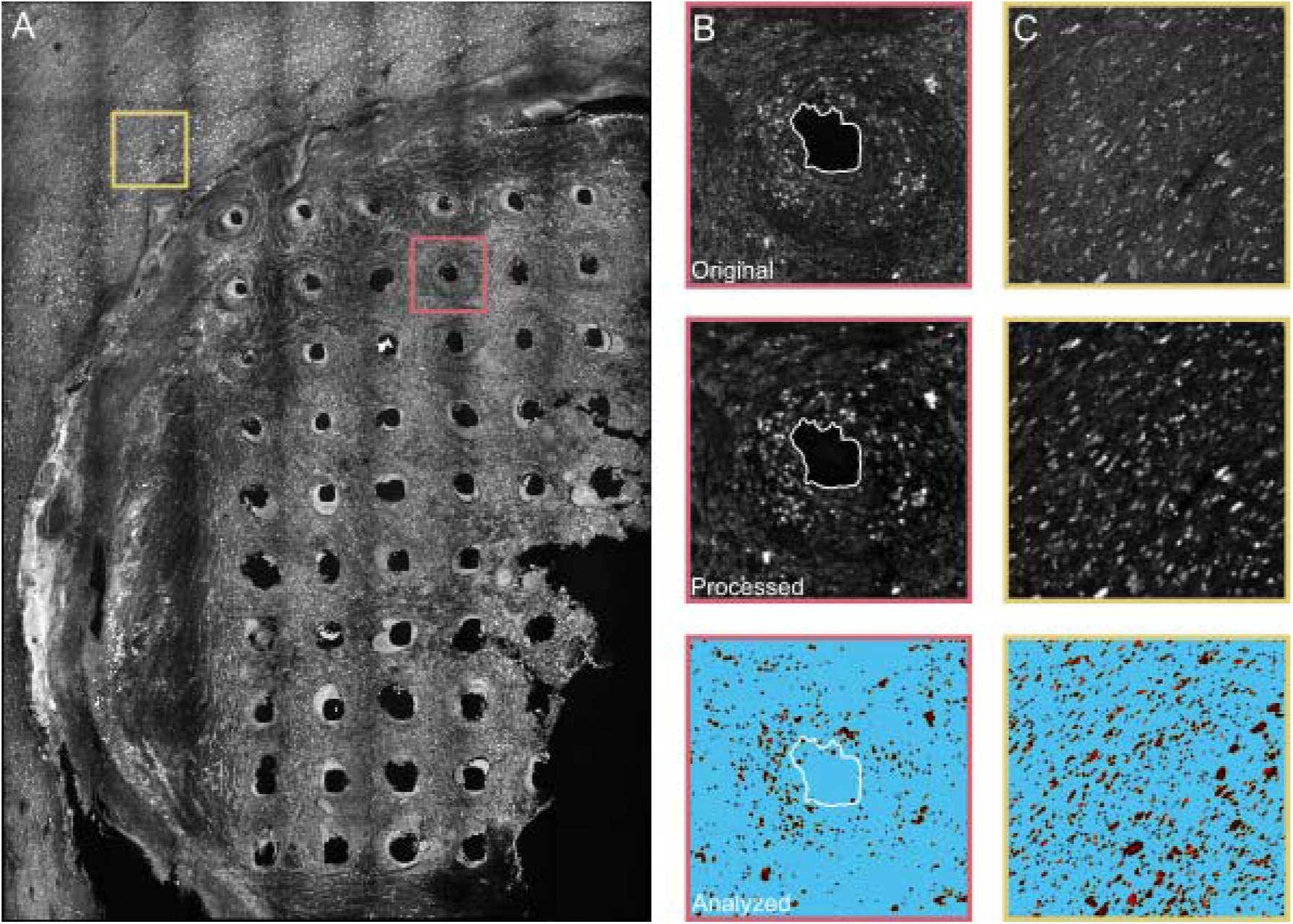
Tissue analysis of NeuN staining of slice 11 in the left sensory cortex under the LS array. A) Confocal image of slice 11 (tissue 1000–1100 µm from brain surface) at a z-stack approximately in the center of the slice. Slice 11 shows 50 intact electrodes holes and nearby non-implanted tissue. The pink box (400 µm x 400 µm) outlines a representative electrode hole seen in (B) and yellow box outlines the non-implanted tissue seen in (C). The representative electrode hole (B) and non-implanted tissue (C) are depicted in original form (top), after filtering and other processing steps (middle), and after analysis in ImageJ with the Analyze Particles program (bottom). Images in (B) and (C) are 400 µm x 400 µm.

For the LS array we determined a non-implanted tissue density of 40.4×10^3^ neurons/mm^3^ for slice 9, 33.8×10^3^ neurons/mm^3^ for slice 11 and 38.2×10^3^ neurons/mm^3^ for slice 13. In comparison, there were fewer neurons in the tissue around the electrode holes. We calculated a mean neuron density surrounding the 393 intact electrodes holes, from the six slices, of 13.9×10^3^ ± 9.6×10^3^ neurons/mm^3^. The neuron density surrounding the electrode holes was reduced by 63% compared to the nearby non-implanted tissue.

### 3.2 Analysis of UEA electrodes

SEM images were collected of the four Utah arrays to identify visible damage or degradation to the electrode shanks. One experimental array (LM) was excluded from the characterization study due to a complete lack of Parylene C and tip coating. We quantified the occurrence of the six degradation categories over the three arrays (N=300 electrode shanks), shown in Figure 3C. When ranked from most to least present, the six categories were Parylene C cracks (40.3%), coating cracks (39.7%), tip breakage (22.3%), shank fracture (3.3%), abnormal debris (1.7%), and Parylene C delamination (1.3%). Of all examined electrodes, 112 electrodes or 37.3% exhibited visible to no degradation.

**Figure 3.**
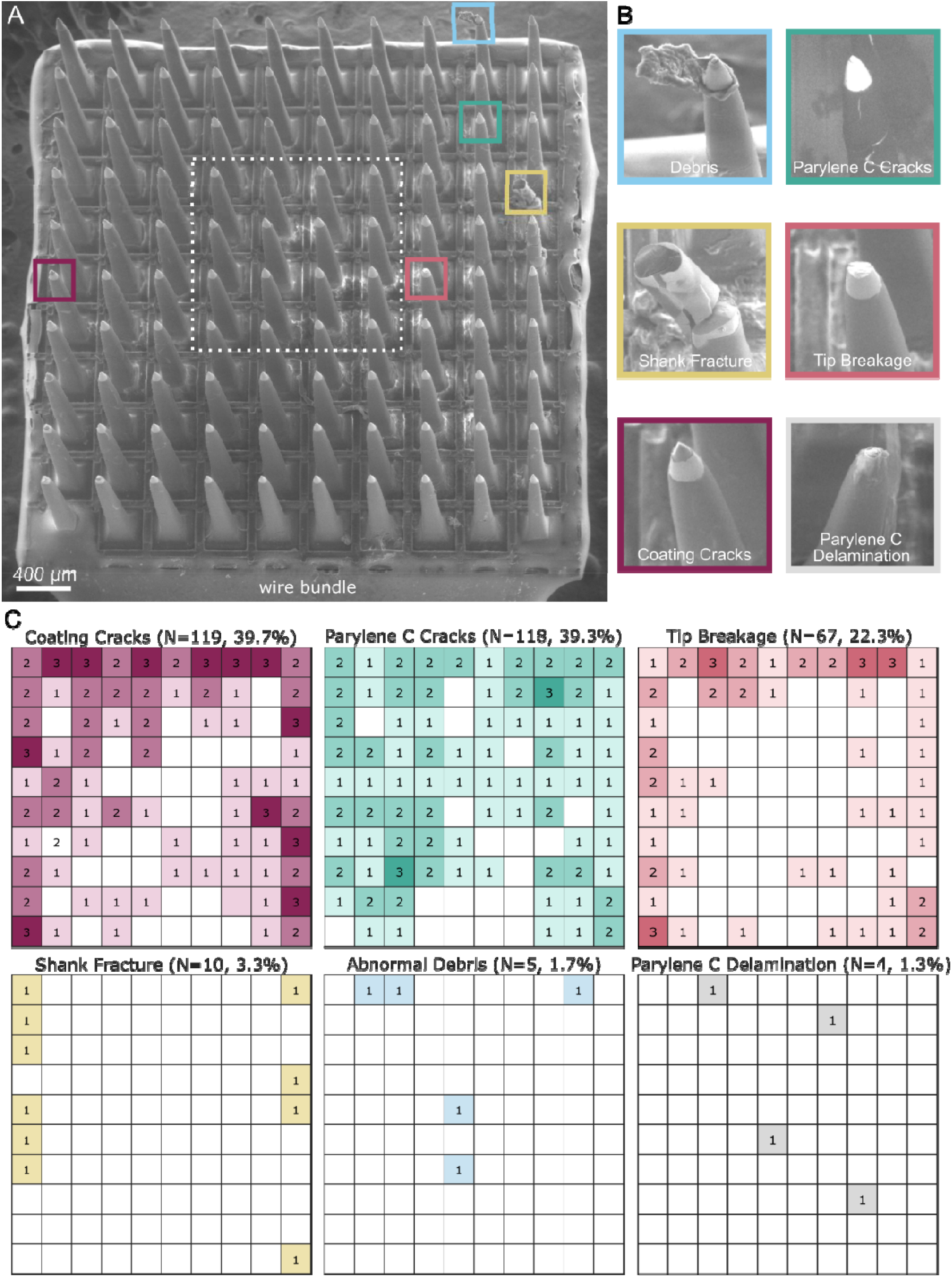
Explanted UEA SEM images and analysis for degradation. A) SEM image of the UEA implanted in the left sensory cortex after 848 days. The UEA is oriented with the wire bundle on the bottom edge. B) Example images of the six categories quantified across three analyzed UEAs. Images of debris, Parylene C cracks, shank fracture, tip breakage, and coating cracks are from the UEA in the left sensory cortex in (A). The representative image of Parylene C delamination is from the UEA in the right motor cortex, as the UEA in the left sensory cortex did not exhibit Parylene C delamination. C) Heat maps of the summation of categorical occurrences in the three analyzed UEAs. The orientation of each heat map is that of the image in (A). Coating cracks occurred most frequently (N=119 electrode shanks, 39.7%), followed by Parylene C cracks (N=118, 39.3%), and then tip breakage (N=67, 22.3%). Coating cracks and tip breakage were significantly more frequent in the outer three rows of electrode shanks than the inner four rows (p-value < 0.05). Shank fracture (N=10, 3.3%), abnormal debris (N=5, 1.7%), and Parylene C delamination (N=4, 1.3%) occurred less frequently.

We analyzed the spatial arrangement of each degradation by performing a 1-way ANOVA test comparing the occurrence of degraded shanks in the outermost three rows of electrode shanks to the innermost 4×4 square of electrode shanks (Figure 3A, dashed white box). There was a significant difference between the outer and inner electrode shanks for the coating cracks (p=0.003) and tip breakage (p=0.004).

## 4. Discussion

In the present study, we explored UEA longevity in the brain through histological analysis of NHP cortex and examination of the mechanical degradation of UEAs implanted for 1.6 and 2.3 years in the brain. The purpose of this study was to enrich our understanding of failure mechanisms in long-term BMIs, which rely on brain signals recorded by UEAs.

Here, we calculated the neuron density surrounding the electrode shank holes and nearby non-implanted tissue. We found a 63% decrease in neuron density surrounding UEA shanks (13.9×10^3^ neurons/mm^3^) compared to that of nearby non-implanted tissue (37.4×10^3^ neurons/mm^3^). Many previous studies have examined the formation of a scar around chronically implanted UEAs [47] but few have quantified the effect UEAs have on the nearby neuron population in NHPs [13], [18], [40], [48], [49]. The neuron density of non-implanted tissue found here is on the same order of magnitude as those in previous studies that found a non-implanted neuron density in the primary motor (66×10^3^ neurons/mm^3^) and primary somatosensory (101×10^3^ neurons/mm^3^) areas of the marmoset NHP cortex [50]. Our preliminary observations note a dramatic decrease in neurons within the immediate recording radius of the electrode, indicating signal loss from long-term UEA usage may be due to a lack of neurons. However, given that the tissue analyzed in this study did not capture the electrode tip, it may be possible that neurons migrated down along the shank to a location where the shank was thinner and perhaps caused less damage.

This study also evaluated SEM images of the extracted UEAs for tip breakage, tip coating cracking, shank fracture, abnormal debris, Parylene C delamination, and Parylene C cracking. Tip breakage, coating cracking, and Parylene C cracking appeared on 20–40% of the electrode shanks, while shank fracture, abnormal debris, and Parylene C delamination appeared on fewer than 4% of shanks. A concentration of degradation at the most vulnerable portion of the electrode shank, the electrode tip, is expected. However, Parylene C cracking would indicate a substantial decrease in electrical impedance, which we did not see from anecdotal evidence. The Parylene C cracks may be a result of the pressure within the SEM, despite being imaged under low vacuum, but this would indicate the silicon shank had detached from the Parylene C coating, another cause for electrical impedance change.

We found a significant difference in the number of electrode shanks exhibiting tip breakage and coating cracks on the outer perimeter of the UEA as compared to the inner region. This may be explained by lateral stresses placed on the outermost electrode shanks when pulled by the wire bundle or micromotion of the brain. Shank fracture, while occurring on just 3% of electrode shanks, may be partially or entirely explained by post-mortem extraction. Our surgical notes indicate that some shank fracture may have occurred during extraction from the fixed tissue, although a precise number is not known.

While this study furthers our knowledge on the impact of chronic UEA implantation, it is also limited. The most obvious limitation is the single NHP used in this study. Higher-order animal models are invaluable to clinical research and studies are constructed to maximize the lifespan and usefulness of each animal. This minimizes access to NHP brain tissue surrounding chronically implanted UEAs [18], [40]. This study was also limited in the number of UEAs implanted in the single NHP. Four UEAs were implanted, one per cortical region of interest in each hemisphere. However, upon termination, one array was discovered to be fully encapsulated in fibrotic tissue. Previous studies have examined the fibrotic tissue response to implanted silicon electrodes and found that fibrotic encapsulation is not an unusual outcome for long term implants in brain [40], [51] or nerve tissue [52]. However, few studies have examined the encapsulated UEA for damage or degradation [31]. This study found that the encapsulated UEA was devoid of metal tip coating material or Parylene C insulation, despite the silicon structure remaining otherwise intact. It is possible that the reactive oxide species in the fibrotic encapsulation caused severe degradation of the Parylene C and metal tip coating, while the fibrotic tissue provided a buffer against physical damage [33].

This study is also limited by the available stained tissue, which constrained the location of non-implant tissue to an area just outside the UEA footprint. While our neuron density of non-implanted tissue aligns with previous records of healthy NHP cortex neuron density [50], the region may still be impacted by the nearby UEA in ways unknown to us [53]. Additionally, the depth of the tissue slices along the length of the electrode shank is not precisely known. We identified the tissue depth as relative to the surface of the brain for that specific tissue section. However, the tissue surface can be irregular or cratered, making it difficult to know the exact depth of the tissue slice. Additionally, we were unable to identify the end of the shank hole within any stained tissue slice. We choose to analyze slices that depicted obvious shank holes, but this limits out ability to know the distance between the recording region and the neurons in the slices analyzed here.

## 5. Conclusion

This study elucidates the effects of UEAs chronically implanted in the motor and sensory cortices of an NHP. While mechanical degradation occurred on 20–40% of electrode shanks, neuronal loss of nearly 63% near the electrode shanks likely contributes more to signal attenuation. Therefore, this work indicates that BMI performance may be more limited by a lack of nearby neurons than material failures of UEAs.

## Acknowledgments

This work was financially supported by the National Institute of General Medical Sciences (R01GM111293), the National Institute of Neurological Disorders and Stroke (UF1NS107659), and the National Science Foundation (1707316).

## Author Contributions

PRP, EJW, HS, AJB, DC, and CAC designed the study. EJW, HS, PRP, and JGL analyzed the histological data. PRP, AVM, and CMC processed, stained, and imaged the tissue. EJW, JMR, and HET, analyzed and scored explanted arrays. PGP performed the NHP surgeries. All authors approved the final manuscript.

## References

[1] J. L. Collinger et al., “High-performance neuroprosthetic control by an individual with tetraplegia,” Lancet, vol. 381, no. 9866, pp. 557–564, 2013.

[2] T. S. Davis et al., “Restoring motor control and sensory feedback in people with upper extremity amputations using arrays of 96 microelectrodes implanted in the median and ulnar nerves,” J. Neural Eng., vol. 13, no. 3, 2016.

[3] D. M. Page et al., “Motor Control and Sensory Feedback Enhance Prosthesis Embodiment and Reduce Phantom Pain After Long-Term Hand Amputation,” Front. Hum. Neurosci., vol. 12, no. 352, 2018.

[4] L. R. Hochberg et al., “Neuronal ensemble control of prosthetic devices by a human with tetraplegia,” Nature, vol. 442, no. 7099, pp. 164–71, 2006.

[5] L. R. Hochberg et al., “Reach and grasp by people with tetraplegia using a neurally controlled robotic arm,” Nature, vol. 485, no. 7398, pp. 372–375, May 2012.

[6] W. Wang, A. D. Degenhart, D. L. Holder, S. Louis, and D. W. Moran, “Human Motor Cortical Activity Recorded with Micro-ECoG Electrodes During Individual Finger Movements,” no. 1, pp. 1–9, 2011.

[7] A. Srikameswaran, “Man With Spinal Cord Injury Uses Brain-Computer Interface to Move Prosthetic Arm With His Thoughts,” PittChronicle, 2011.

[8] A. B. Ajiboye et al., “Restoration of reaching and grasping movements through brain-controlled muscle stimulation in a person with tetraplegia: a proof-of-concept demonstration,” Lancet, vol. 389, no. 10081, pp. 1821–1830, 2017.

[9] S. N. Flesher et al., “Intracortical microstimulation of human somatosensory cortex,” Sci. Transl. Med., vol. 8, no. 361, pp. 1–10, 2016.

[10] P. Souza, “President Obama fist-bumps Nathan Copeland,” Obama White House, 2016. [Online]. Available: https://obamawhitehouse.archives.gov/photos-and-video/photo/2016/10/president-obama-fist-bumps-nathan-copeland.

[11] K. A. Ludwig, R. M. Miriani, N. B. Langhals, M. D. Joseph, D. J. Anderson, and D. R. Kipke, “Using a common average reference to improve cortical neuron recordings from microelectrode arrays,” J. Neurophysiol., vol. 101, no. 3, pp. 1679–1689, 2009.

[12] S. R. Nason et al., “A low-power band of neuronal spiking activity dominated by local single units improves the performance of brain–machine interfaces,” Nat. Biomed. Eng., vol. 4, no. 10, pp. 973–983, 2020.

[13] P. J. Rousche and R. A. Normann, “Chronic recording capability of the utah intracortical electrode array in cat sensory cortex,” J. Neurosci. Methods, vol. 82, no. 1, pp. 1–15, 1998.

[14] R. A. Normann, E. M. Maynard, P. J. Rousche, and D. J. Warren, “A neural interface for a cortical vision prosthesis,” Vision Res., vol. 39, no. 15, pp. 2577–2587, 1999.

[15] C. T. Nordhausen, E. M. Maynard, and R. A. Normann, “Single unit recording capabilities of a 100 microelectrode array,” Brain Res., vol. 726, no. 1–2, pp. 129–140, 1996.

[16] R. A. Normann and E. Fernandez, “Clinical applications of penetrating neural interfaces and Utah Electrode Array technologies,” J. Neural Eng., vol. 13, no. 6, 2016.

[17] N. Y. Masse et al., “Non-causal spike filtering improves decoding of movement intention for intracortical BCIs,” J. Neurosci. Methods, vol. 236, pp. 58–67, 2014.

[18] J. C. Barrese et al., “Failure mode analysis of silicon-based intracortical microelectrode arrays in non-human primates,” J. Neural Eng., vol. 10, no. 6, 2013.

[19] J. D. Simeral, S. P. Kim, M. J. Black, J. P. Donoghue, and L. R. Hochberg, “Neural control of cursor trajectory and click by a human with tetraplegia 1000 days after implant of an intracortical microelectrode array,” J. Neural Eng., vol. 8, no. 2, 2011.

[20] A. J. Bullard, B. C. Hutchison, J. Lee, C. A. Chestek, and P. G. Patil, “Estimating Risk for Future Intracranial, Fully Implanted, Modular Neuroprosthetic Systems: A Systematic Review of Hardware Complications in Clinical Deep Brain Stimulation and Experimental Human Intracortical Arrays,” Neuromodulation, vol. 23, no. 4, pp. 411–426, 2020.

[21] C. Sponheim et al., “Longevity and reliability of chronic unit recordings using the Utah, intracortical multi-electrode arrays,” J. Neural Eng., vol. 18, no. 6, 2021.

[22] R. Biran, D. C. Martin, and P. A. Tresco, “The brain tissue response to implanted silicon microelectrode arrays is increased when the device is tethered to the skull,” J. Biomed. Mater. Res. - Part A, vol. 82, no. 1, pp. 169–178, 2007.

[23] L. Karumbaiah et al., “Relationship between intracortical electrode design and chronic recording function,” Biomaterials, vol. 34, no. 33, pp. 8061–8074, Nov. 2013.

[24] T. D. Y. Kozai et al., “Mechanical failure modes of chronically implanted planar silicon-based neural probes for laminar recording,” Biomaterials, vol. 37, pp. 25–39, Jan. 2015.

[25] J. W. Salatino, K. A. Ludwig, T. D. Y. Kozai, and E. K. Purcell, “Glial responses to implanted electrodes in the brain,” Nat. Biomed. Eng., vol. 1, no. 11, pp. 862–877, 2017.

[26] K. A. Potter, A. C. Buck, W. K. Self, and J. R. Capadona, “Stab injury and device implantation within the brain results in inversely multiphasic neuroinflammatory and neurodegenerative responses,” J. Neural Eng., vol. 9, no. 4, 2012.

[27] N. F. Nolta, M. B. Christensen, P. D. Crane, J. L. Skousen, and P. A. Tresco, “BBB leakage, astrogliosis, and tissue loss correlate with silicon microelectrode array recording performance,” Biomaterials, vol. 53, pp. 753–762, 2015.

[28] B. J. Black et al., “Chronic recording and electrochemical performance of utah microelectrode arrays implanted in rat motor cortex,” J. Neurophysiol., vol. 120, no. 4, pp. 2083–2090, 2018.

[29] R. Biran, D. C. Martin, and P. A. Tresco, “Neuronal cell loss accompanies the brain tissue response to chronically implanted silicon microelectrode arrays,” Exp. Neurol., vol. 195, no. 1, pp. 115–126, 2005.

[30] D. A. Henze, Z. Borhegyi, J. Csicsvari, A. Mamiya, K. D. Harris, and G. Buzsáki, “Intracellular features predicted by extracellular recordings in the hippocampus in vivo,” J. Neurophysiol., vol. 84, no. 1, pp. 390–400, 2000.

[31] R. Caldwell, M. G. Street, R. Sharma, P. Takmakov, B. Baker, and L. Rieth, “Characterization of Parylene-C degradation mechanisms: In vitro reactive accelerated aging model compared to multiyear in vivo implantation,” Biomaterials, vol. 232, no. December 2019, p. 119731, 2020.

[32] S. F. Cogan, “Neural Stimulation and Recording Electrodes,” Annu. Rev. Biomed. Eng., vol. 10, no. 1, pp. 275–309, 2008.

[33] E. S. Ereifej et al., “Implantation of neural probes in the brain elicits oxidative stress,”Front. Bioeng. Biotechnol., vol. 6, no. FEB, 2018.

[34] B. Shafer, C. Welle, and S. Vasudevan, “A rat model for assessing the long-term safety and performance of peripheral nerve electrode arrays,” J. Neurosci. Methods, vol. 328, no. August, p. 108437, 2019.

[35] D. H. Szarowski et al., “Brain responses to micro-machined silicon devices,” Brain Res., vol. 983, no. 1–2, pp. 23–35, 2003.

[36] M. P. Ward, P. Rajdev, C. Ellison, and P. P. Irazoqui, “Toward a comparison of microelectrodes for acute and chronic recordings,” Brain Res., vol. 1282, pp. 183–200, Jul. 2009.

[37] D. McCreery, S. Cogan, S. Kane, and V. Pikov, “Correlations between histology and neuronal activity recorded by microelectrodes implanted chronically in the cerebral cortex,” J. Neural Eng., vol. 13, no. 3, 2016.

[38] K. Woeppel et al., “Explant Analysis of Utah Electrode Arrays Implanted in Human Cortex for Brain-Computer-Interfaces,” Front. Bioeng. Biotechnol., vol. 9, no. December, pp. 1–15, 2021.

[39] L. J. Szymanski et al., “Neuropathological effects of chronically implanted, intracortical microelectrodes in a tetraplegic patient,” J. Neural Eng., vol. 18, no. 4, 2021.

[40] J. C. Barrese, J. Aceros, and J. P. Donoghue, “Scanning electron microscopy of chronically implanted intracortical microelectrode arrays in non-human primates,” J. Neural Eng., vol. 13, no. 2, 2016.

[41] A. J. Bullard, “Feasibility of Using the Utah Array for Long-Term Fully Implantable Neuroprosthesis Systems,” University of Michigan, 2019.

[42] Z. T. Irwin et al., “Neural control of finger movement via intracortical brain-machine interface,” J. Neural Eng., vol. 14, no. 6, 2017.

[43] A. K. Vaskov et al., “Cortical Decoding of Individual Finger Group Motions Using ReFIT Kalman Filter,” Front. Neurosci., vol. 12, no. November, 2018.

[44] K. E. Schroeder et al., “Disruption of corticocortical information transfer during ketamine anesthesia in the primate brain,” Neuroimage, vol. 134, pp. 459–465, 2016.

[45] X. Xie et al., “Bi-layer encapsulation of utah array based nerual interfaces by atomic layer deposited Al2O3 and parylene C,” 2013 Transducers Eurosensors XXVII 17th Int. Conf. Solid-State Sensors, Actuators Microsystems, TRANSDUCERS EUROSENSORS 2013, no. June, pp. 1267–1270, 2013.

[46] K. Miura, “Bleach correction ImageJ plugin for compensating the photobleaching of time-lapse sequences,” F1000Research, vol. 9, pp. 1–17, 2020.

[47] M. B. Christensen, S. M. Pearce, N. M. Ledbetter, D. J. Warren, G. A. Clark, and P. A. Tresco, “The foreign body response to the Utah Slant Electrode Array in the cat sciatic nerve,” Acta Biomater., vol. 10, no. 11, 2014.

[48] K. A. Malaga et al., “Data-driven model comparing the effects of glial scarring and interface interactions on chronic neural recordings in non-human primates,” J. Neural Eng., vol. 13, no. 1, 2016.

[49] P. A. House, J. D. MacDonald, P. A. Tresco, and R. A. Normann, “Acute microelectrode array implantation into human neocortex: preliminary technique and histological considerations.,” Neurosurg. Focus, vol. 20, no. 5, pp. 19–22, 2006.

[50] N. Atapour, P. Majka, I. H. Wolkowicz, D. Malamanova, K. H. Worthy, and M. G. P. Rosa, “Neuronal Distribution across the Cerebral Cortex of the Marmoset Monkey (Callithrix jacchus),” Cereb. Cortex, vol. 29, no. 9, pp. 3836–3863, 2019.

[51] P. A. Cody, J. R. Eles, C. F. Lagenaur, T. D. Y. Kozai, and X. T. Cui, “Unique electrophysiological and impedance signatures between encapsulation types: An analysis of biological Utah array failure and benefit of a biomimetic coating in a rat model,” Biomaterials, vol. 161, pp. 117–128, 2018.

[52] A. Branner, R. B. Stein, E. Fernandez, Y. Aoyagi, and R. A. Normann, “Long-Term Stimulation and Recording with a Penetrating Microelectrode Array in Cat Sciatic Nerve,” IEEE Trans. Biomed. Eng., vol. 51, no. 1, pp. 146–157, 2004.

[53] C. H. Thompson, A. Saxena, N. Heelan, J. Salatino, and E. K. Purcell, “Spatiotemporal patterns of gene expression around implanted silicon electrode arrays,” J. Neural Eng., vol. 18, no. 4, 2021.

